# Semi-automated Production of Cell-free Biosensors

**DOI:** 10.1101/2024.10.13.618078

**Authors:** Dylan M. Brown, Daniel A. Phillips, David C. Garcia, Anibal Arce, Tyler Lucci, John P. Davies, Jacob Mangini, Katherine A. Rhea, Casey B. Bernhards, John R. Biondo, Steven M. Blum, Stephanie D. Cole, Jennifer A. Lee, Marilyn S. Lee, Nathan D. McDonald, Brenda Wang, Dale L. Perdue, Walter Thavarajah, Ashty S. Karim, Matthew W. Lux, Michael C. Jewett, Aleksandr E. Miklos, Julius B. Lucks

**Affiliations:** Department of Chemical and Biological Engineering, Northwestern University, Evanston, IL 60208, USA; Center for Synthetic Biology, Northwestern University, Evanston, IL 60208, USA; Applied Synthetic Biology and Olfaction, U.S. Army DEVCOM Chemical Biological Center, Gunpowder, MD 21010, USA; Precise Systems, Lexington Park, MD 20653, USA; Decontamination Sciences Branch, U.S. Army DEVCOM Chemical Biological Center. Gunpowder MD, 21010, USA; University of Maryland, College Park, MD 20742, USA; Defense Threat Reduction Agency, Fort Belvoir, VA 22060, USA; Biomanufacturing Branch, U.S. Army DEVCOM Chemical Biological Center, Gunpowder, MD 21010, USA; Department of Bioengineering, Stanford University, Stanford, CA 94305, USA

**Keywords:** Cell-free biosensors, cell-free systems, point-of-use manufacturing, fluoride riboswitch, automation

## Abstract

Cell-free synthetic biology biosensors have potential as effective *in vitro* diagnostic technologies for the detection of chemical compounds such as toxins and human health biomarkers. They have several advantages over conventional laboratory-based diagnostic approaches, including being able to be assembled, freeze-dried, distributed, and then used at the point-of-need. This makes them an attractive platform for cheap and rapid chemical detection across the globe. Though promising, a major challenge is scaling up biosensor manufacturing to meet the needs of their multiple uses. Currently, cell-free biosensor assembly during lab-scale development is mostly performed manually by the operator, leading to quality control and performance variability issues. Here we explore the use of liquid handling robotics to manufacture cell-free biosensor reactions. We compare both manual and semi-automated reaction assembly approaches using the Opentrons OT-2 liquid handling platform on two different cell-free gene expression assay systems that constitutively produce colorimetric (LacZ) or fluorescent (GFP) signals. We test the designed protocol by constructing an entire 384-well plate of fluoride sensing cell-free biosensors and demonstrate that they perform closely to expected detection outcomes.

## Introduction

Environmental chemical hazards are a major global threat to human and environmental health – affecting air, water, soil, and food quality globally.^1–5^ Exposure to environmental hazards leads to poor human health outcomes such as asthma, mental illness, birth defects, cancer, chronic illness, cardiovascular disease or death.^6–10^ Current technologies for assessing environmental hazards often require laboratory facilities, electronic devices, and technical expertise. Though a variety of government organizations monitor contaminants, their methods of detection often include analytical techniques such as gas chromatography, liquid chromatography, and mass spectroscopy.^11,12^ Field deployable technologies are prone to similar user-difficulties, requiring expertise in the field or complex data analysis, with modern methods still using electrical signals, chromatography, or spectroscopic devices.^13–16^ These types of detection methods are often inaccessible to under-resourced locations and difficult to use for individuals without experience who would benefit most from easy-to-use detection methods. Additionally, these detection methods lack scalability, in that detection reliant on standard devices is often not rapid and cannot be mass produced and distributed to individuals easily and cheaply.

Cell-free gene expression systems can be used as a powerful strategy to cheaply create ready-to-use diagnostic devices that are able to be freeze-dried and easily deployed at the point-of-need.^17– 19^These sensors take advantage of cellular machinery for detection of small molecules and ions. Additionally, these reactions are relatively shelf stable disposable, and biodegradable.^20–25^ Cell-free biosensors have demonstrated functionality in field deployment applications, being used to detect copper in water from California, fluoride levels in water from Kenya and Costa Rica, and used for educational purposes in high schools.^26–29^

In each of these studies, cell-free biosensors were manufactured manually, and sensor quality was variable, motivating the need to develop approaches that can produce easy-to-use field deployable sensors with consistent quality at scale. This is particularly important in cases where rapid generation of these diagnostics are needed. Automated approaches that incorporate robotic liquid handlers have the potential to allow for higher numbers of sensors to be manufactured with expected quality consistency across production batches.

Here we sought to adapt these approaches to manufacture cell-free biosensors process to a robotic toolkit. We specifically chose the Opentrons OT-2 device as a platform that is widely used across biology to improve experimental workflows, with the added advantage of its affordability when compared to other robotics systems. Additionally, Opentrons devices have a low barrier to entry with lower expertise requirement and the potential to be easily deployed for site specific manufacturing. To investigate the use of the Opentrons OT-2 system for biosensor manufacturing, we first characterized non-optimized robotic production compared to manual production means, and then developed an automated protocol to produce hundreds of biosensor reactions, which were confirmed to be functional with follow-on characterization. We anticipate that this strategy can be widely applied to other biosensor systems and general cell-free system applications, such as cell-free protein synthesis, or cell-free metabolic engineering, and can be further scaled to meet application needs.

## Results

Automation can be used to improve reproducibility and standardize processes within synthetic biology.^30^ Automation workflows can be designed under a variety of regimes, where the method of reaction construction can greatly affect manufacturing outcomes. The first step of determining automation best practices is the selection of manufacturing regimes and factors that are best suited for process outcomes.^31–33^ To determine this, we used two separate modes of reaction preparation: (i) manual reaction assembly and (ii) automated assembly with an Opentrons OT-2 liquid handling robot. In addition, we explored two modes of assembly: individual mix and master mix approaches where the individual mix approach describes the process of transferring each reaction component (DNA, cell extract, and reaction buffer components) separately to their respective tubes and the master mix configuration uses a premixed reaction, where the DNA, cell extract, and reaction buffer is mixed then distributed into the reaction tubes. Individual mixtures may allow for more flexibility of the reaction environment and allow for component variation, while the master mix approach is quick and easier to carry out in bulk for reactions that are compositionally consistent. For a small set of reactions created by a trained experimenter, master and individual mix approaches perform similarly, however for “out of the box” robotic approaches the individual mix method leads to more failures, whereas the master mix approach is closer to human performance (**Supplemental Figure 1**).

This led us to adopt the master mix approach, as biosensor reactions for a given target can be produced with a bulk mixture given the desire to create reactions with homogenous compositions. To characterize robot performance at a larger scale before optimization, we assessed constitutive cell-free reaction performance over 288 manually constructed reactions (the total of 96 reactions carried out by 3 different experimenters) and 288 automation constructed reactions (the total of 96 reactions carried out by 3 different experimenters) for two different reporter systems, totaling 1,152 reactions (**Figure 1**). We employed different experimenters to assess if robotic performance could address variability in reaction performance imparted by manual construction by multiple individuals. Reaction master mixes were manually created and then either placed on the Opentrons robot for distribution or distributed by hand. Reactions were then lyophilized for 16 hours and characterized the next day post rehydration (**Figure 1A**). To compare manual versus pre-optimized robotic reactions we used two metrics indicative of reaction performance; maximum signal and the time to reach half of the maximum signal (t_1/2_) (**Figure 1B**).

**Figure 1.**
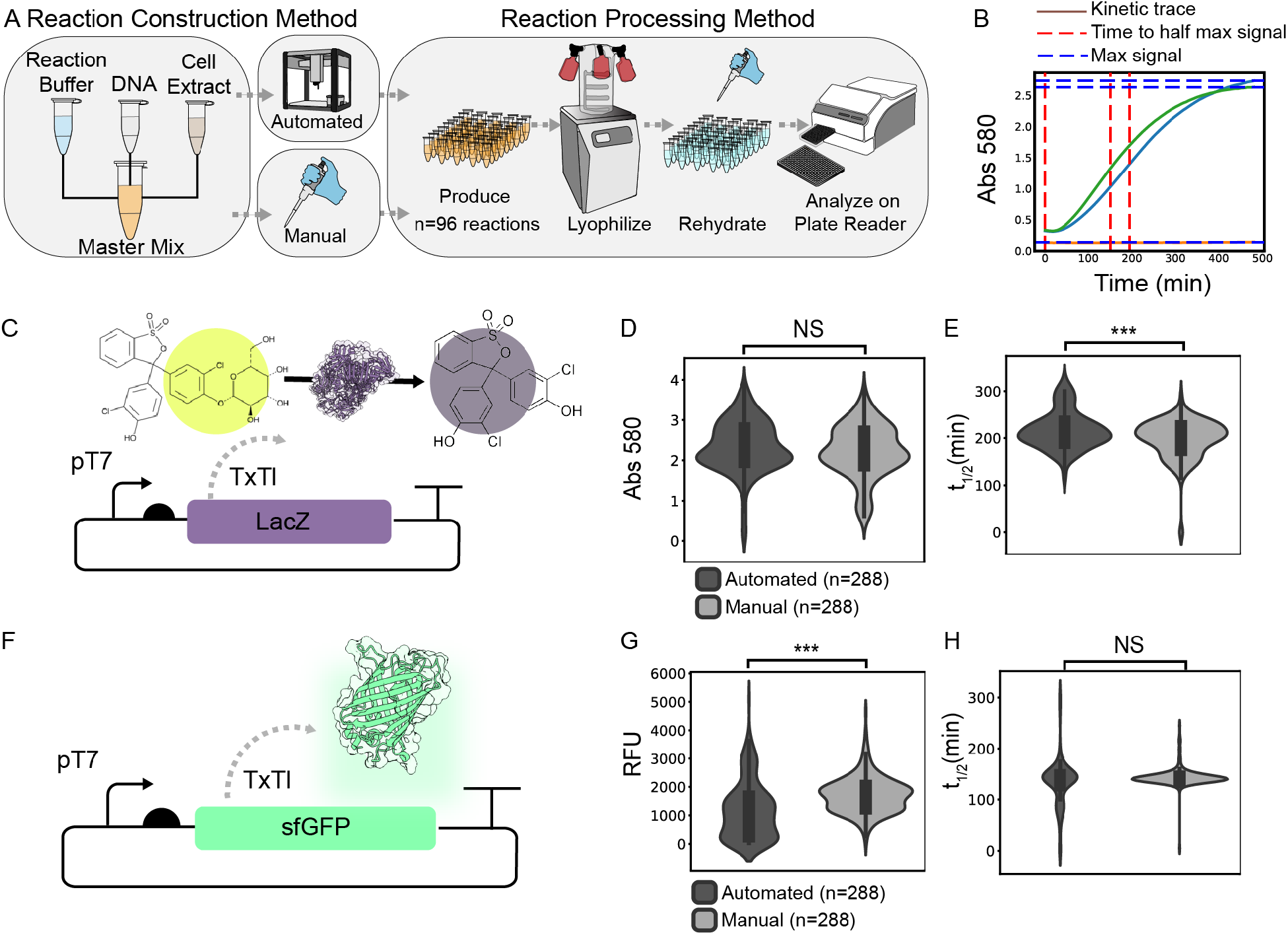
Characterization of manual and robotic construction of cell-free reactions. A) Schematic of master mix reaction construction methods for manual and automated approaches. B) Representative example of data processing method, where t_1/2_ and max signal are calculated and stored for each kinetic curve (two shown) in a cell-free reaction population. C) Schematic of constitutive LacZ expression producing a colorimetric signal. Violin plots showing D) maximum colorimetric signal, or E) t_1/2_ values from constitutive LacZ producing reactions for n=288 manual and n=288 automated reactions. For each set of 288 reactions, 3 different experimenters constructed 96 of these reactions. F) Schematic of constitutive T7 GFP expression producing a fluorescent signal. G) Violin plots showing G) max fluorescent signal or H) t_1/2_ values from constitutive GFP producing reactions for n=288 manual reactions and n=288 automated reactions. For each set of 288 reactions, 3 different experimenters constructed 96 of these reactions.

To test the effects of different reporter systems, we assembled cell-free reactions that either constitutively expressed a LacZ enzymatic reporter that produces a colorimetric signal, or super-folder green fluorescent protein (sfGFP) reporter that produces a fluorescence signal (**Figure 1C, 1F**). For the enzymatic reporter, non-optimized automated protocols showed significant differences in t_1/2_ when compared across manual and automated construction approaches with non-significant differences to maximum signal (**Figure 1D, 1E**). Alternatively, for the fluorescent protein reporter we see that t_1/2_ characterization shows no significant differences between automated and manual approaches, however, the variance in the maximum signal was significantly different between these conditions (**Figure 1G, 1H**). Overall, this shows that kinetic properties were retained well for a fluorescent reporter, where variations mostly occur for maximum signal response metrics, while for enzymatic reporters the opposite was true perhaps due to the dye concentration thresholding the maximal signal that can be generated by these reactions. This demonstrates differences in robotic and human performance for different reporter systems. Additionally, robotic systems appear unable to improve consistency imparted by multiple experimenters using basal robot settings.

Because of the inconsistency in reaction performance for robot constructed reactions, we developed a protocol to improve both reaction quality and reduce the burden of manual scale-up (**Figure 2A**). Because the reactions are viscous, we found that a variety of parameters have effects on the distribution efficacy, including the dispense and aspirate rates, number of mixes, volume of mixing, liquid blowout height, touch-tip height, and dispense heights. Blowout refers to the process of fully removing excess liquid from the pipette after dispensing, while touch tip touches the sides of the reaction vessel walls post blowout to remove excess liquid that may remain on the tip. We observed that without modifying these robotic parameters, the reaction easily bubbled and got stuck in the pipette tips causing aspiration errors for subsequent reactions. To help remediate this issue, we included further tip changes to prevent accumulation of reaction in the tips. We found that the combination of lower dispensing heights and dispense rate, blowing out excess cell-free expression (CFE) mix, and using the touch tip function allowed for better distribution.

**Figure 2.**
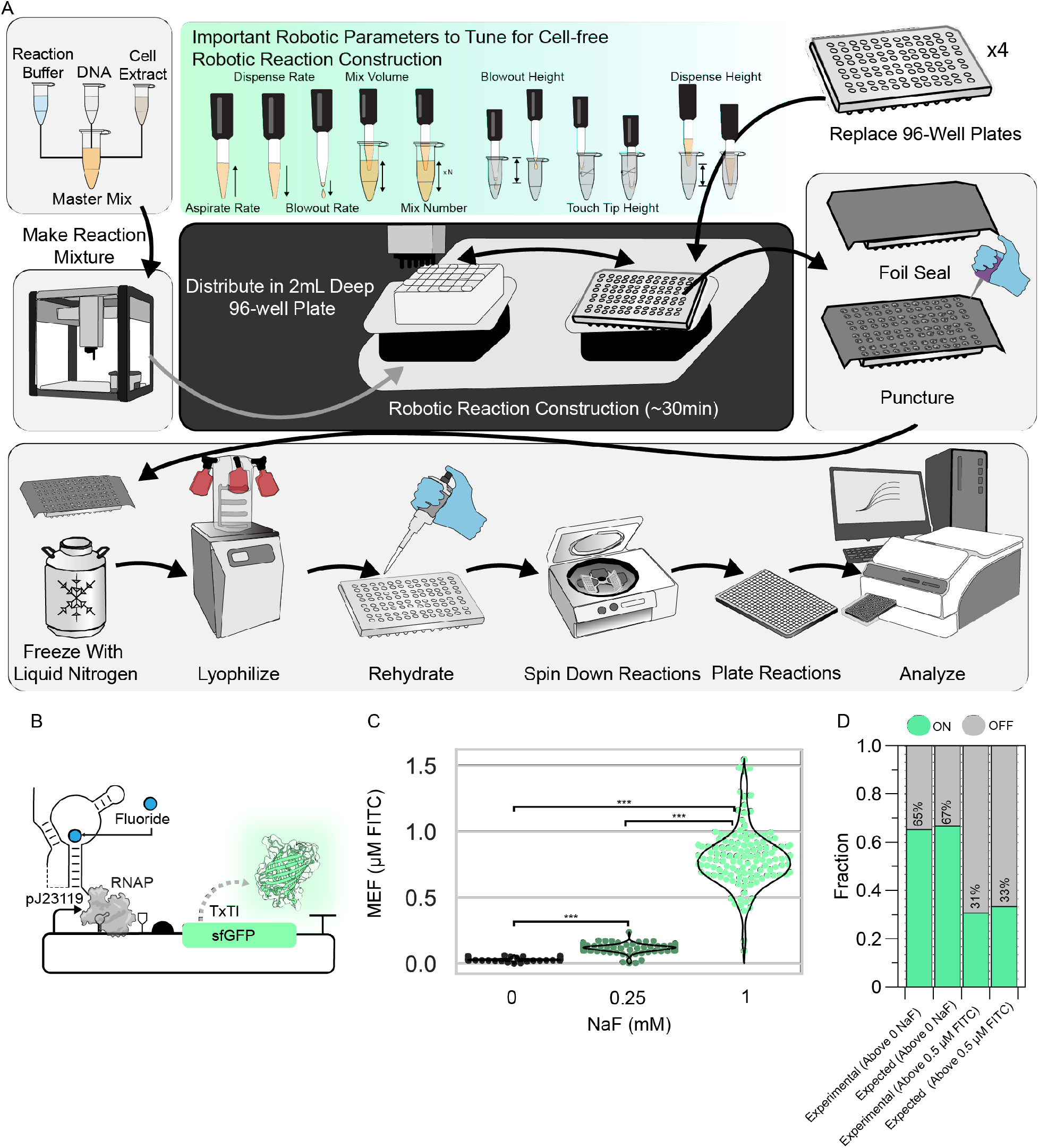
Characterization of robotic performance for constructing cell-free fluoride riboswitch biosensors. A) Schematic of semi-automated workflow for constructing hundreds of biosensor reactions using the Opentrons robotic device. B) Schematic of fluoride riboswitch biosensor mechanism. C) Violin plot of maximum signal for n=384 biosensor reactions. Three concentrations of fluoride were used to hydrate the reactions distributed amongst the plates using an interleaved-signal format to account for distribution effects. D) Classification of reaction performance compared to expected performance using two different performance criteria. The first criteria represent the fraction of reactions that are expected to have a signal value higher than the maximum signal of the zero NaF condition which are the 0.25 nM and 1 mM NaF conditions. The second criteria are used for biosensor field deployment metrics, where fluorescence can be visualized with a hand-held device when signal is above 0.5 uM FITC. Here it is expected that only the 1mM NaF meets this criterion.

To test this protocol, we constructed 384 fluoride riboswitch reactions and characterized subpopulations of these reactions with zero, 0.25 mM and 1 mM NaF concentrations. The protocol can construct this number of reactions in approximately 30 minutes, with the user swapping out 96-well PCR plates. This substantially reduces the experimenter burden on constructing the reactions. The fluoride riboswitch populations were then assessed based on maximum signal post lyophilization. All reactions were successfully dispensed in this process. Here, we can see sensor populations follow a more consistent distribution pattern when compared to pre-optimized distributions (**Figure 2C**). Additional analysis shows that 65% of the total reaction population achieved an ON state (fluorescence above the 0mM NaF condition) compared to 67% for expected ON states which was the percentage of reactions that contained either 0.25 mM or 1 mM NaF (**Figure 2D**). Previous work has shown that a fluorescence level of 0.5 μM FITC calibrated signal is visualizable by eye with a detection device.^27^ We therefore investigated the number of reactions that met this condition (**Figure 2D**). The expected value of signal > 0.5 μM FITC was calculated as the number of samples exposed to 1mM NaF over the total reaction population, while the experimental data is the fraction of reactions that achieved this. Here we observed 31% of reactions met this threshold compared to the expected value of 33%. Overall, the observed reaction performance was in line with expected performance in the reactions distributed by the Opentrons OT-2 device.

## Discussion

There is a need to easily and accessibly scale-up the production of cell-free systems such as cell-free biosensors for point-of-use applications. Here we present an approach that addresses this need by integrating easy-to-use robotics protocols to automate the assembly of reactions that can be freeze dried in bulk before use by simple rehydration. We demonstrate this scale-up using the fluoride riboswitch, which has been previously deployed for point-of-use studies in Kenya and Costa Rica.^28,29^, showing that our approach can assemble hundreds of reactions that perform as expected.

The tools described in this study can be adapted to a variety cell-free reaction regimes, with the protocol designed to distribute reactions for 12 96-well plates at a time before needing to be restarted. This allows for producing upwards of thousands of reactions, likely needing few modifications to the approach depending on the Opentrons device and experimenter needs. As applications of cell-free manufacturing and biosensing become more realized, the ability to produce this scale of reactions in bench top settings or lower-resourced environments becomes enabling.^34–37^ A particular selling point of the Opentrons device is that it can be easily relocated and started up, which furthers applications for accessible point-of-use diagnostics production. This automated approach also reduces potential user error from manual methods that can have a large effect on batch-to-batch variation, and rather leads to consistent and expected experimental population behavior for the fluoride riboswitch^38,39^.

To optimize the system, we needed to assess parameters that effect robotic performance when pipetting viscous liquids. As such, mixing rates, liquid blowout height, tip-touch height, and swapping out multichannel pipette tips were all considered as factors that can be adjusted to reduce reaction failures and misfiring. It is important to consider these aspects when porting the protocol to other systems, and these parameters may be tuned depending on Opentrons device differences, software versions, and reaction compositions. Additionally, it should be noted that rehydration procedures can also influence reaction outcomes as it involves pipetting steps that can be subject to variability.

To facilitate adoption of this approach we provide detailed protocols and Opentrons operation code that implements the procedures described here. Our goal is for this protocol to serve as an accessible starting point to produce cell-free reactions at scales that match many application needs.

## Materials and Methods

### Plasmids

For pT7 sfGFP synthesis, pJL1 (Addgene #69496) was used. For pY71-LacZ gene expression a new plasmid was constructed. pJBL3752 (Addgene #128809) was used for fluoride detection. Sequences for plasmids can be found in **Supplemental Information Table 1**.

### E. coli Lysate Production

Lysate preparation was carried out using E. coli BL21DE3* for extract used in the fluoride riboswitch experiments and Rosetta (DE3) pLysS *ΔlacZα* strains for the analysis in Figure 1. Preparation for E. coli BL21(DE3)* extract was carried out using methods previously published.^40^ Briefly, BL21(DE3)* cells were plated on agar and a single colony was cultured overnight then inoculated into 1 L of 2X YT + P media, composed of 16g tryptone, 10g yeast extract, 5g NaCl, 7g potassium phosphate dibasic, 3g potassium phosphate monobasic. Cells were then grown with shaking at 220 rpm at 37°C and harvested at OD600 3.0, after approximately 4 hours. Once an OD600 of 3.0 was reached, the cells were processed in accordance with the previous protocol up through dialysis.^40^ A 100L culture of Rosetta (DE3) *ΔlacZ* cells was processed for lysate production similar to the BL21(DE3)* lysate production mentioned previously, but with modifications to accommodate production at scale. Briefly, 750 mL starter cultures (1.5 L total) were grown for 16 hours at 37°C with 200 rpm shaking incubation. Prior to inoculating 100 L of 2X YT + P culture media supplemented with 5 mL of antifoam 204 (Sigma, A8311) in an IF 150L (New Brunswick Scientific) fermenter, the media was allowed to aerate overnight with a rotor speed of 100 rpm and 20 standard liters per minute (slpm) airflow at 37°C. After inoculation with enough overnight culture to yield a starting OD600 of 0.05, the fermenter settings were adjusted to 300 rpm, 50 splm, and the dissolved oxygen (DO) was calibrated to 100%. At an OD600 of 0.6-1.0, the culture was induced with a final concentration of 1 mM isopropyl B-D-1-thiogalatopyranoside (IPTG) (GoldBio, I2481C). Once DO reached 50%, the rotor speed was increased to 500 rpm. At an OD600 of 3.5, the culture was cooled to 4°C, centrifuged in a prechilled Powerfuge pilot, 1.1 L bowl system (CARR Biosystems) within approximately 8 hours, and the pelleted bacteria was subsequently processed as described previously.^40^

### Constitutive Cell-free Reaction Assay

Reaction master mix for pT7-sfGFP and pY71-LacZ constitutive expression was carried out using a Cytomim master mix optimized by Cai et al., which contains 8mM magnesium glutamate, 260mM potassium glutamate, 1.26mM AMP, 0.86mM GMP, 0.86mM UMP, 0.86mM CMP, 4mM oxalic acid, 2mM L-glutathione, 1.5mM spermidine, 9.2mM potassium phosphate dibasic, 5.8mM potassium phosphate monobasic, 2mM amino acids, and 1mM tyrosine.^41^ Eleven μL reactions were assembled using 4.4 μL Cai master mix, with 3.3 μL cell extract, and 3.3 μL water with DNA concentrated to 15nM in the complete reaction. For LacZ reporter reactions, 12.5mM chlorophenol red b-D-galactopyranoside (CPRG) was also added into the water component of the reaction to a final volume of 0.88 μL (1mM concentration). Cell-free reactions were set up for 96, 11μL reactions, with 20% dead volume for a total of 1267μL volume scaled proportionally to the volumes of individual reaction components listed above.

Cell-free expression reactions were carried out manually or using default Opentrons OT-2 settings. For manual expression, a master mix containing the above components was made and hand mixed 20 times using a p1000 pipette set to half the total volume (633μL). The master mix was then distributed into 11μL aliquots using a p20 pipette into 96 PCR tubes on ice. For robotic distributions, the master mix was made by hand without mixing and then placed on an Opentrons 24 1.5mL tube rack. The robot was then made to mix the reaction 20 times at default rates using the Opentrons p1000 pipette attachment. The robot then distributed 11μL of this master mix into 96 PCR tubes on a cooling module set to 4°C using a p20 pipette attachment.

This process was completed by 3 separate experimenters, using the same manual and automated protocol (**Supplemental Data and Code**). Once reactions were completed, they were immediately frozen with liquid nitrogen and placed on a Labconco FreeZone 2.5 Liter -84C Benchtop Freeze Dryer for 16 hours. Immediately after lyophilization, reactions were rehydrated with 11μL of water using a multichannel pipette, spun down, mixed 20 times by hand, and plated on a 384 well plate. CPRG reporter reactions were then read at absorbance 585 nm wavelength at 5-minute intervals for 480 minutes. Fluorescent reporter reactions were monitored using excitation/emission 485nm/520nm at 5-minute intervals for 480 mins. All reactions were analyzed at 30°C.

### Fluoride Riboswitch Cell-free Reaction Assay

Cell-free reactions for the fluoride riboswitch were carried out using a modified phosphoenolpyruvate, amino acids, NAD+, and oxalic acid (PANOx)^42^ reaction system with salt solution containing 8mM magnesium glutamate, 10 mM ammonium glutamate, 130 mM potassium glutamate; transcription master mix with 1.2 mM ATP, 0.850 mM GTP, 0.850 mM UTP, 0.850 mM CTP, 72μM folinic acid, 0.171 mg/mL tRNA; amino acids solution with 2 mM amino acids; energy solution of 30 mM PEP; and cofactor solution with 0.33 mM NAD, 0.27 mM CoA, 4 mM oxalic acid, 1 mM putrescine, 1.5 mM spermidine, 57 mM HEPES.^40^ Eleven μL reactions were assembled using 3.3 μL PANOx master mix, with 3.3 μL cell extract, and 4.4 μL fluoride riboswitch DNA diluted in water to a final reaction concentration of 15nM. The reactions were constructed using the master mix approach, where a bulk solution of 6083 μL was generated which includes a 20% dead volume adjustment for robotic procedures. This reaction was then manually distributed in the first row of a 2mL 96 deep-well blocks with 760μL of reaction in each well. This 96-well block was then placed on the Opentrons OT-2 platform for use by the protocol provided in **Supplemental Data and Code**. The Opentrons OT-2 platform configuration is shown in **Supplemental Information Figure 2**. The reactions were distributed in 11μL aliquots into 96-well PCR plates and kept on ice post distribution until all plates were completed, for a total of 4 plates. These plates were then simultaneously immersed in liquid nitrogen on 96-well aluminum PCR tube blocks and immediately placed on a Labconco FreeZone 2.5 Liter -84C Benchtop Freeze Dryer for 16 hours using a 4 shelf lyophilization chamber. Immediately after lyophilization, the reactions were manually rehydrated with a multichannel pipette the next day with the indicated concentrations of NaF using an interleaved-signal format to account for positional effects further described in **Supplemental Information Figure 3**. The reactions were then monitored using excitation/emission 485nm/520nm at 5-minute intervals for 10 hours at 30 °C.

### Data analysis code and statistics

JMP Pro 16 was used to carry out non-parametric statistical analysis for data gathered in Figure 1 and Figure 2. Additionally, Python code for data analysis is included in in **Supplemental Data and Code** with instructional and layout information on how to structure and carry out Opentrons protocols found in Figure 2 (**Supplemental Information Figure 2, 3**). Opentrons protocols are available and annotated in supplemental information.

## Data Availability

All data presented are available alongside code for analysis and Opentrons operation at https://github.com/LucksLab/Brown_Phillips_Semi-Automated_Biosensor_Manufacturing_2024/tree/main.

## Code Availability

Python code for data analysis as well as Opentrons protocols can be found at https://github.com/LucksLab/Brown_Phillips_Semi-Automated_Biosensor_Manufacturing_2024/tree/main. This contains excel and python files containing the raw data from Figure 1 and Figure 2 as well as output files and image files generated from the code. All code for data analysis was written with the assistance of OpenAI ChatGPT 3.5 and manually edited.

## Author Contributions

D.M.B., J.B.L., D.A.P., D.C.G., A.E.M. designed the study. D.M.B, T.J.L., A.A.M., D.A.P. carried out experimental work in the manuscript. D.M.B., J.M., D.A.P, D.C.G. developed Opentrons protocols used in the study. D.M.B developed analytical code. J.P.D., D.A.P., D.C.G, J.M. determined important robotic system parameters for tuning. K.A.R, J.R.B. generated materials and lysate scale-up procedures used within. M.L., W.T., C.B.B., S.M.B., S.D.C, J.A.L., N.D.M., B.W., D.L.P., aided in experimentation and procedure development. J.B.L., A.M., M.C.J., A.S.K. supervised the study and acquired funding. D.M.B., D.A.P., J.B.L., and A.E.M. wrote the manuscript. All authors edited the manuscript.

## Acknowledgements

This work was also supported by Army Contracting Command (W52P1J-21-9-3023), and the National Science Foundation (2310382) to J.B.L. J.B.L. was also supported by a John Simon Guggenheim Fellowship. The views, opinions, and/or findings expressed are those of the authors and should not be interpreted as representing the official views or policies of the Department of Defense, the National Science Foundation or the U.S. Government.

## References

1. Carvalho, F. P. Pesticides, environment, and food safety. Food Energy Secur. 6, 48–60 (2017).

2. Carvalho, F. P. Agriculture, pesticides, food security and food safety. Environ. Sci. Policy 9, 685–692 (2006).

3. Md Meftaul, I., Venkateswarlu, K., Dharmarajan, R., Annamalai, P. & Megharaj, M. Pesticides in the urban environment: A potential threat that knocks at the door. Sci. Total Environ. 711, 134612 (2020).

4. Rizzo, D. M., Lichtveld, M., Mazet, J. A. K., Togami, E. & Miller, S. A. Plant health and its effects on food safety and security in a One Health framework: four case studies. One Health Outlook 2021 31 3, 1–9 (2021).

5. WHO. Guidelines for drinking-water quality, 4th edition, incorporating the 1st addendum. (2017).

6. Human Exposure and Health | US EPA. https://www.epa.gov/report-environment/human-exposure-and-health.

7. Eales, J. et al. Human health impacts of exposure to phthalate plasticizers: An overview of reviews. Environ. Int. 158, 106903 (2022).

8. Sharma, N. & Singhvi, R. Effects of Chemical Fertilizers and Pesticides on Human Health and Environment: A Review. Int. J. Agric. Environ. Biotechnol. 10, 675 (2017).

9. Fuller, R. et al. Pollution and health: a progress update. Lancet Planet. Health 6, e535–e547 (2022).

10. Rovira, J. & Domingo, J. L. Human health risks due to exposure to inorganic and organic chemicals from textiles: A review. Environ. Res. 168, 62–69 (2019).

11. Chartres, N., Bero, L. A. & Norris, S. L. A review of methods used for hazard identification and risk assessment of environmental hazards. Environ. Int. 123, 231–239 (2019).

12. Collection of Methods | US EPA. https://www.epa.gov/measurements-modeling/collection-methods#3.

13. Kaserzon, S. L., Heffernan, A. L., Thompson, K., Mueller, J. F. & Gomez Ramos, M. J. Rapid screening and identification of chemical hazards in surface and drinking water using high resolution mass spectrometry and a case-control filter. Chemosphere 182, 656–664 (2017).

14. Schierenbeck, T. M. & Smith, M. C. Path to Impact for Autonomous Field Deployable Chemical Sensors: A Case Study of in Situ Nitrite Sensors. Environ. Sci. Technol. 51, 4755– 4771 (2017).

15. Benasco, A. R. et al. Receptor Induced Doping of Conjugated Polymer Transistors: A Strategy for Selective and Ultrasensitive Phosphate Detection in Complex Aqueous Environments. Adv. Electron. Mater. 8, 2101353 (2022).

16. Clark, R. B. & Dick, J. E. Towards deployable electrochemical sensors for per- and polyfluoroalkyl substances (PFAS). Chem. Commun. 57, 8121–8130 (2021).

17. Thavarajah, W. et al. A primer on emerging field-deployable synthetic biology tools for global water quality monitoring. Npj Clean Water 3, 1–10 (2020).

18. Carlson, E. D., Gan, R., Hodgman, C. E. & Jewett, M. C. Cell-free protein synthesis: Applications come of age. Biotechnol. Adv. 30, 1185–1194 (2012).

19. Silverman, A. D., Karim, A. S. & Jewett, M. C. Cell-free gene expression: an expanded repertoire of applications. Nat. Rev. Genet. 21, 151–170 (2020).

20. Arce, A. et al. Decentralizing Cell-Free RNA Sensing With the Use of Low-Cost Cell Extracts. Front. Bioeng. Biotechnol. 9, (2021).

21. Warfel, K. F. et al. A Low-Cost, Thermostable, Cell-Free Protein Synthesis Platform for On-Demand Production of Conjugate Vaccines. ACS Synth. Biol. 12, 95–107 (2023).

22. Stark, J. C. et al. On-demand biomanufacturing of protective conjugate vaccines. Sci. Adv. 7, eabe9444 (2021).

23. Pardee, K. et al. Paper-Based Synthetic Gene Networks. Cell 159, 940–954 (2014).

24. Pardee, K. et al. Portable, On-Demand Biomolecular Manufacturing. Cell 167, 248-259.e12 (2016).

25. Pardee, K. et al. Rapid, Low-Cost Detection of Zika Virus Using Programmable Biomolecular Components. Cell 165, 1255–1266 (2016).

26. Jung, K. J. et al. At-home, cell-free synthetic biology education modules for transcriptional regulation and environmental water quality monitoring. bioRxiv 2023.01.09.523248 (2023) doi:10.1101/2023.01.09.523248.

27. Jung, J. K. et al. Cell-free biosensors for rapid detection of water contaminants. Nat. Biotechnol. 2020 3812 38, 1451–1459 (2020).

28. Thavarajah, W. et al. Point-of-Use Detection of Environmental Fluoride via a Cell-Free Riboswitch-Based Biosensor. ACS Synth. Biol. 9, 10–18 (2020).

29. Thavarajah, W. et al. The accuracy and usability of point-of-use fluoride biosensors in rural Kenya. Npj Clean Water 2023 61 6, 1–8 (2023).

30. Jessop-Fabre, M. M. & Sonnenschein, N. Improving reproducibility in synthetic biology. Front. Bioeng. Biotechnol. 7, 18 (2019).

31. Hofmann, P., Samp, C. & Urbach, N. Robotic process automation. Electron. Mark. 30, 99– 106 (2020).

32. Syed, R. et al. Robotic Process Automation: Contemporary themes and challenges. Comput. Ind. 115, 103162 (2020).

33. Herm, L. V. et al. A framework for implementing robotic process automation projects. Inf. Syst. E-Bus. Manag. 1–35 (2022) doi:10.1007/S10257-022-00553-8/FIGURES/3.

34. Thames, A. H. et al. A Cell-Free Protein Synthesis Platform to Produce a Clinically Relevant Allergen Panel. ACS Synth. Biol. 12, 2252–2261 (2023).

35. DeWinter, M. A. et al. Point-of-Care Peptide Hormone Production Enabled by Cell-Free Protein Synthesis. ACS Synth. Biol. 12, 1216–1226 (2023).

36. Ji, X.Liu, W.-Q. & Li, J. Recent advances in applying cell-free systems for high-value and complex natural product biosynthesis. Curr. Opin. Microbiol. 67, 102142 (2022).

37. Hu, V. T. & Kamat, N. P. Cell-free protein synthesis systems for vaccine design and production. Curr. Opin. Biotechnol. 79, 102888 (2023).

38. Cole, S. D. et al. Quantification of Interlaboratory Cell-Free Protein Synthesis Variability. ACS Synth. Biol. 8, 2080–2091 (2019).

39. Rhea, K. A. et al. Variability in cell-free expression reactions can impact qualitative genetic circuit characterization. Synth. Biol. 7, ysac011 (2022).

40. Silverman, A. D., Kelley-Loughnane, N., Lucks, J. B. & Jewett, M. C. Deconstructing Cell-Free Extract Preparation for in Vitro Activation of Transcriptional Genetic Circuitry. ACS Synth. Biol. 8, 403–414 (2019).

41. Cai, Q. et al. A simplified and robust protocol for immunoglobulin expression in Escherichia coli cell-free protein synthesis systems. Biotechnol. Prog. 31, 823–831 (2015).

42. Kim, D.-M. & Swartz, J. R. Regeneration of adenosine triphosphate from glycolytic intermediates for cell-free protein synthesis. Biotechnol. Bioeng. 74, 309–316 (2001).

