## Supplementary Information for "Semi-automated Production of Cell-free Biosensors"

**Supplemental Table 1.** Plasmid sequences

| Plasmid | Sequence |
|---|---|
| pT7-sfGFP (pJL1 Addgene #69496)<br><br>sfGFP<br>KanR<br>ColE1 Origin of Replication<br>T7 Promoter<br>RBS<br>T7 Terminator | agatcaaaggatcttcttgagatcctttttttctgcgcgtaatctgctgcttgcaaacaaaaaaaccaccgctaccag<br>cggtggtttgtttgccggatcaagagctaccaactctttttccgaaggtaactggcttcagcagagcgcagatacc<br>aaatactgttcttctagtgtagccgtagttaggccaccacttcaagaactctgtagcaccgcctacatacctcgctct<br>gctaatcctgttaccagtggctgctgccagtggcgataagtcgtgtcttaccgggttggactcaagacgatagttac<br>cggataaggcgcagcggtcgggctgaacggggggttcgtgcacacagcccagcttggagcgaacgacctac<br>accgaactgagatacctacagcgtgagctatgagaaagcgccacgcttcccgaagggagaaaggcggacagg<br>tatccggtaagcggcagggtcggaacaggagagcgcacgagggagcttccagggggaaacgcctggtatctt<br>tatagtcctgtcgggtttcgccacctctgacttgagcgtcgatttttgtgatgctcgtcaggggggcggagcctatg<br>gaaaaacgccagcaacgcgatcccgcgaaattaatacgactcactataggggagaccacaacggtttccctctag<br>aaataattttgtttaactttaagaaggagatatacatATGAGCAAAGGTGAAGAACTGTTTA<br>CCGGCGTTGTGCCGATTCTGGTGGAACTGGATGGCGATGTGAACGG<br>TCACAAATTCAGCGTGCGTGGTGAAGGTGAAGGCGATGCCACGATT<br>GGCAAACTGACGCTGAAATTTATCTGCACCACCGGCAAACTGCCGG<br>TGCCGTGGCCGACGCTGGTGACCACCCTGACCTATGGCGTTCAGTGT<br>TTTAGTCGCTATCCGGATCACATGAAACGTCACGATTTCTTTAAATC<br>TGCAATGCCGGAAGGCTATGTGCAGGAACGTACGATTAGCTTTAAA<br>GATGATGGCAAATATAAAACGCGCGCCGTTGTGAAATTTGAAGGCG<br>ATACCCTGGTGAACCGCATTGAACTGAAAGGCACGGATTTTAAAGA<br>AGATGGCAATATCCTGGGCCATAAACTGGAATACAACTTTAATAGC<br>CATAATGTTTATATTACGGCGGATAAACAGAAAAATGGCATCAAAG<br>CGAATTTTACCGTTCGCCATAACGTTGAAGATGGCAGTGTGCAGCTG<br>GCAGATCATTATCAGCAGAATACCCCGATTGGTGATGGTCCGGTGC<br>TGCTGCCGGATAATCATTATCTGAGCACGCAGACCGTTCTGTCTAAA<br>GATCCGAACGAAAAAGGCACGCGGGACCACATGGTTCTGCACGAAT<br>ATGTGAATGCGGCAGGTATTACGTGGAGCCATCCGCAGTTCGAAAA<br>ATAAgtcgaccggctgctaacaaagcccgaaaggaagctgagttggctgctgccaccgctgagcaataact<br>agcataaccccttggggcctctaaacgggtcttgaggggttttttgctgaaagccaattctgattagaaaaactcat |

| | |
|---|---|
| | cgagcatcaaatgaaactgcaatttattcatatcaggattatcaataccatatttttgaaaaagccgtttctgtaatgaa<br>ggagaaaactcaccgaggcagttccataggatggcaagatcctggtatcggtctgcgattccgactcgtccaaca<br>tcaatacaacctattaatttcccctcgtcaaaaataaggttatcaagtgagaaatcaccatgagtgacgactgaatcc<br>ggtgagaatggcaaaagcttatgcatttctttccagacttgttcaacaggccagccattacgctcgtcatcaaaatc<br>actcgcatcaaccaaaccgttattcattcgtgattgcgcctgagcgagacgaaatacgcgatcgctgttaaaagga<br>caattacaaacaggaatcgaatgcaaccggcgcaggaacactgccagcgcatcaacaatattttcacctgaatca<br>ggatattcttctaatacctggaatgctgttttcccggggatcgcagtggtgagtaaccatgcatcatcaggagtacg<br>gataaaatgcttgatggtcggaagaggcataaattccgtcagccagtttagtctgaccatctcatctgtaacatcatt<br>ggcaacgctacctttgccatgtttcagaaacaactctggcgcatcgggcttcccatacaatcgatagattgtcgcac<br>ctgattgcccgacattatcgcgagcccatttatacccatataaatcagcatccatgttggaatttaatcgcggcttcg<br>agcaagacgtttcccgttgaatatggctcataacacccccttgtattactgtttatgtaagcagacagttttattgttcat<br>gatgatatatttttatcttgtgcaatgtaacatcagagattttgagacacaacgtg |
| pY71-LacZ<br><br>LacZ<br>KanR<br>ColE1 Origin of Replication<br>T7 Promoter<br>sTRSV HHRz<br>RBS | TAAtaaacaataactgaatagggatcccgactggcgagagccaggtaacgaatggatccttcagccaaaaaa<br>cttaagaccgccggtcttgtccactaccttgcagtaatgcggtggacaggatcggcggttttcttttctcttctcaag<br>ctgaaagccaattctgattagaaaaactcatcgagcatcaaatgaaactgcaatttattcatatcaggattatcaata<br>ccatatttttgaaaaagccgtttctgtaatgaaggagaaaactcaccgaggcagttccataggatggcaagatcct<br>ggtatcggtctgcgattccgactcgtccaacatcaatacaacctattaatttcccctcgtcaaaaataaggttatcaa<br>gtgagaaatcaccatgagtgacgactgaatccggtgagaatggcaaaagcttatgcatttctttccagacttgttca<br>acaggccagccattacgctcgtcatcaaaatcactcgcatcaaccaaaccgttattcattcgtgattgcgcctgagc<br>gagacgaaatacgcgatcgctgttaaaaggacaattacaaacaggaatcgaatgcaaccggcgcaggaacact<br>gccagcgcatcaacaatattttcacctgaatcaggatattcttctaatacctggaatgctgttttcccggggatcgca<br>gtggtgagtaaccatgcatcatcaggagtacggataaaatgcttgatggtcggaagaggcataaattccgtcagc<br>cagtttagtctgaccatctcatctgtaacatcattggcaacgctacctttgccatgtttcagaaacaactctggcgcat<br>cgggcttcccatacaatcgatagattgtcgcacctgattgcccgacattatcgcgagcccatttatacccatataaat<br>cagcatccatgttggaatttaatcgcggcttcgagcaagacgtttcccgttgaatatggctcataacacccccttgtat<br>tactgtttatgtaagcagacagttttattgttcatgatgatatatttttatcttgtgcaatgtaacatcagagattttgagac<br>acaacgtggatcctgcagttgagatcctttttttctgcgcgtaatctgctgcttgcaaacaaaaaaaccaccgctacc<br>agcggtggtttgtttgccggatcaagagctaccaactctttttccgaaggtaactggcttcagcagagcgcagata<br>ccaaatactgtccttctagtgtagccgtagttaggccaccacttcaagaactctgtagcaccgcctacatacctcgc<br>tctgctaatcctgttaccagtggctgctgccagtggcgataagtcgtgtcttaccgggttggactcaagacgatagt<br>taccggataaggcgcagcggtcgggctgaacggggggttcgtgcacacagcccagcttggagcgaacgacct<br>acaccgaactgagatacctacagcgtgagcattgagaaagcgccacgcttcccgaagggagaaaggcggaca<br>ggtatccggtaagcggcagggtcggaacaggagagcgcacgagggagcttccagggggaaacgcctggtat<br>ctttatagtcctgtcgggtttcgccacctctgacttgagcgtcgatttttgtgatgctcgtcaggggggcggagccta<br>tggaaacgaattcagatctcgatcccgcgaaattaatacgactcactatagggagctgtcaccggatgtgctttcc<br>ggtctgatgagtccgtgaggacgaaacagcctctacaaataattttgtttaatctagagaaagaggagaaatacta<br>gATGACCATGATTACGGATTCACTGGCCGTCGTTTTACAACGTCGTG<br>ACTGGGAAAACCCTGGCGTTACCCAACTTAATCGCCTTGCAGCACA<br>TCCCCCTTTCGCCAGCTGGCGTAATAGCGAAGAGGCCCGCACCGAT<br>CGCCCTTCCCAACAGTTGCGCAGCCTGAATGGCGAATGGCGCTTTGC<br>CTGGTTTCCGGCACCAGAAGCGGTGCCGGAAAGCTGGCTGGAGTGC<br>GATCTTCCTGAGGCCGATACTGTCGTCGTCCCCTCAAACTGGCAGAT<br>GCACGGTTACGATGCGCCCATCTACACCAACGTGACCTATCCCATTA<br>CGGTCAATCCGCCGTTTGTTCCCACGGAGAATCCGACGGGTTGTTAC<br>TCGCTCACATTTAATGTTGATGAAAGCTGGCTACAGGAAGGCCAGA<br>CGCGAATTATTTTTGATGGCGTTAACTCGGCGTTTCATCTGTGGTGC<br>AACGGGCGCTGGGTCGGTTACGGCCAGGACAGTCGTTTGCCGTCTG<br>AATTTGACCTGAGCGCATTTTTACGCGCCGGAGAAAACCGCCTCGC<br>GGTGATGGTGCTGCGCTGGAGTGACGGCAGTTATCTGGAAGATCAG<br>GATATGTGGCGGATGAGCGGCATTTTCCGTGACGTCTCGTTGCTGCA<br>TAAACCGACTACACAAATCAGCGATTTCCATGTTGCCACTCGCTTTA<br>ATGATGATTTCAGCCGCGCTGTACTGGAGGCTGAAGTTCAGATGTG<br>CGGCGAGTTGCGTGACTACCTACGGGTAACAGTTTCTTTATGGCAGG<br>GTGAAACGCAGGTCGCCAGCGGCACCGCGCCTTTCGGCGGTGAAAT<br>TATCGATGAGCGTGGTGGTTATGCCGATCGCGTCACACTACGTCTGA<br>ACGTCGAAAACCCGAAACTGTGGAGCGCCGAAATCCCGAATCTCTA<br>TCGTGCGGTGGTTGAACTGCACACCGCCGACGGCACGCTGATTGAA<br>GCAGAAGCCTGCGATGTCGGTTTCCGCGAGGTGCGGATTGAAAATG<br>GTCTGCTGCTGCTGAACGGCAAGCCGTTGCTGATTCGAGGCGTTAAC<br>CGTCACGAGCATCATCCTCTGCATGGTCAGGTCATGGATGAGCAGA<br>CGATGGTGCAGGATATCCTGCTGATGAAGCAGAACAACTTTAACGC<br>CGTGCGCTGTTCGCATTATCCGAACCATCCGCTGTGGTACACGCTGT<br>GCGACCGCTACGGCCTGTATGTGGTGGATGAAGCCAATATTGAAAC<br>CCACGGCATGGTGCCAATGAATCGTCTGACCGATGATCCGCGCTGG<br>CTACCGGCGATGAGCGAACGCGTAACGCGAATGGTGCAGCGCGATC<br>GTAATCACCCGAGTGTGATCATCTGGTCGCTGGGGAATGAATCAGG<br>CCACGGCGCTAATCACGACGCGCTGTATCGCTGGATCAAATCTGTC<br>GATCCTTCCCGCCCGGTGCAGTATGAAGGCGGCGGAGCCGACACCA<br>CGGCCACCGATATTATTTGCCCGATGTACGCGCGCGTGGATGAAGA<br>CCAGCCCTTCCCGGCTGTGCCGAAATGGTCCATCAAAAAATGGCTTT |

| | |
|---|---|
| | CGCTACCTGGAGAGACGCGCCCGCTGATCCTTTGCGAATACGCCCA<br>CGCGATGGGTAACAGTCTTGGCGGTTTCGCTAAATACTGGCAGGCG<br>TTTCGTCAGTATCCCCGTTTACAGGGCGGCTTCGTCTGGGACTGGGT<br>GGATCAGTCGCTGATTAAATATGATGAAAACGGCAACCCGTGGTCG<br>GCTTACGGCGGTGATTTTGGCGATACGCCGAACGATCGCCAGTTCTG<br>TATGAACGGTCTGGTCTTTGCCGACCGCACGCCGCATCCAGCGCTGA<br>CGGAAGCAAAACACCAGCAGCAGTTTTTCCAGTTCCGTTTATCCGG<br>GCAAACCATCGAAGTGACCAGCGAATACCTGTTCCGTCATAGCGAT<br>AACGAGCTCCTGCACTGGATGGTGGCGCTGGATGGTAAGCCGCTGG<br>CAAGCGGTGAAGTGCCTCTGGATGTCGCTCCACAAGGTAAACAGTT<br>GATTGAACTGCCTGAACTACCGCAGCCGGAGAGCGCCGGGCAACTC<br>TGGCTCACAGTACGCGTAGTGCAACCGAACGCGACCGCATGGTCAG<br>AAGCCGGGCACATCAGCGCCTGGCAGCAGTGGCGTCTGGCGGAAAA<br>CCTCAGTGTGACGCTCCCCGCCGCGTCCCACGCCATCCCGCATCTGA<br>CCACCAGCGAAATGGATTTTTGCATCGAGCTGGGTAATAAGCGTTG<br>GCAATTTAACCGCCAGTCAGGCTTTCTTTCACAGATGTGGATTGGCG<br>ATAAAAAACAACTGCTGACGCCGCTGCGCGATCAGTTCACCCGTGC<br>ACCGCTGGATAACGACATTGGCGTAAGTGAAGCGACCCGCATTGAC<br>CCTAACGCCTGGGTCGAACGCTGGAAGGCGGCGGGCCATTACCAGG<br>CCGAAGCAGCGTTGTTGCAGTGCACGGCAGATACACTTGCTGATGC<br>GGTGCTGATTACGACCGCTCACGCGTGGCAGCATCAGGGGAAAACC<br>TTATTTATCAGCCGGAAAACCTACCGGATTGATGGTAGTGGTCAAAT<br>GGCGATTACCGTTGATGTTGAAGTGGCGAGCGATACACCGCATCCG<br>GCGCGGATTGGCCTGAACTGCCAGCTGGCGCAGGTAGCAGAGCGGG<br>TAAACTGGCTCGGATTAGGGCCGCAAGAAAACTATCCCGACCGCCT<br>TACTGCCGCCTGTTTTGACCGCTGGGATCTGCCATTGTCAGACATGT<br>ATACCCCGTACGTCTTCCCGAGCGAAAACGGTCTGCGCTGCGGGAC<br>GCGCGAATTGAATTATGGCCCACACCAGTGGCGCGGCGACTTCCAG<br>TTCAACATCAGCCGCTACAGTCAACAGCAACTGATGGAAACCAGCC<br>ATCGCCATCTGCTGCACGCGGAAGAAGGCACATGGCTGAATATCGA<br>CGGTTTCCATATGGGGATTGGTGGCGACGACTCCTGGAGCCCGTCA<br>GTATCGGCGGAATTCCAGCTGAGCGCCGGTCGCTACCATTACCAGTT<br>GGTCTGGTGTCAAAAA |
| pJBL3752 (Addgene #128809)<br><br>J23119 (SpeI) Promoter<br>Fluoride Riboswitch<br>RBS<br>sfGFP<br>p15a Origin of Replication<br>CamR<br>rrnB T1 Terminator | ggagatttcctggaagatgccaggaagatacttaacagggaagtgagagggccgcggcaaagccgttttccat<br>aggctccgcccccctgacaagcatcacgaaatctgacgctcaaatcagtggtggcgaaacccgacaggactat<br>aaagataccaggcgtttccccctggcggctccctcgtgcgctctcctgttcctgcctttcggtttaccggtgtcattc<br>cgctgttatggccgcgtttgtctcattccacgcctgacactcagttccgggtaggcagttcgctccaagctggactg<br>tatgcacgaaccccccgttcagtccgaccgctgcgccttatccggtaactatcgtcttgagtccaacccggaaag<br>acatgcaaaagcaccactggcagcagccactggtaattgatttagaggagttagtcttgaagtcatgcgccggtta<br>aggctaaactgaaaggacaagttttggtgactgcgctcctccaagccagttacctcggttcaaagagttggtagct<br>cagagaaccttcgaaaaaccgccctgcaaggcggttttttcgttttcagagcaagagattacgcgcagaccaaaa<br>cgatctcaagaagatcatcttattaatcagataaaatatttctagatttcagtgcaatttatctcttcaaatgtagcacct<br>gaagtcagccccatacgatataagttgtaattctcatgtttgacagcttatcatcgataagcttccgatggcgcgccg<br>agaggctttacactttatgcttccggctgaattctaaagatcttttgacagctagctcagtcctaggtataatactagttt<br>ataggcgatggagttcgccataaacgctgcttagctaatgactcctaccagtatcactactggtaggagtctattttttt<br>taggaggaaggatctATGAGCAAAGGAGAAGAACTTTTCACTGGAGTTGTCC<br>CAATTCTTGTTGAATTAGATGGTGATGTTAATGGGCACAAATTTTCT<br>GTCCGTGGAGAGGGTGAAGGTGATGCTACAAACGGAAAACTCACCC<br>TTAAATTTATTTGCACTACTGGAAAACTACCTGTTCCGTGGCCAACA<br>CTTGTCACTACTCTGACCTATGGTGTTCAATGCTTTTCCCGTTATCCG<br>GATCACATGAAACGGCATGACTTTTTCAAGAGTGCCATGCCCGAAG<br>GTTATGTACAGGAACGCACTATATCTTTCAAAGATGACGGGACCTA<br>CAAGACGCGTGCTGAAGTCAAGTTTGAAGGTGATACCCTTGTTAAT<br>CGTATCGAGTTAAAGGGTATTGATTTTAAAGAAGATGGAAACATTC<br>TTGGACACAAACTCGAGTACAACTTTAACTCACACAATGTATACATC<br>ACGGCAGACAAACAAAAGAATGGAATCAAAGCTAACTTCAAAATTC<br>GCCACAACGTTGAAGATGGTTCCGTTCAACTAGCAGACCATTATCA<br>ACAAAATACTCCAATTGGCGATGGCCCTGTCCTTTTACCAGACAACC<br>ATTACCTGTCGACACAATCTGTCCTTTCGAAAGATCCCAACGAAAA<br>GCGTGACCACATGGTCCTTCTTGAGTTTGTAACTGCTGCTGGGATTA<br>CACATGGCATGGATGAGCTCTACAAATAAggatccaaactcgagtaaggatctccag<br>gcatcaaataaaacgaaaggctcagtcgaaagactgggcctttcgttttatctgttgtttgtcggtgaacgctctcta<br>ctagagtcacactggctcaccttcgggtgggcctttctgcgtttatacctagggtacggttttgggatccttactcga<br>gtctagactgcagttgatcgggcacgtaagaggttccaactttcaccataatgaaataagatcactaccgggcgta<br>ttttttgagttatcgagattttcaggagctaaggaagctaaaatggagaaaaaaatcactggatataccaccgttgat<br>atatcccaatggcatcgtaaagaacattttgaggcatttcagtcagttgctcaatgtacctataaccagaccgttcag<br>ctggatattacggcctttttaaagaccgtaaagaaaaataagcacaagttttatccggcctttattcacattcttgccc<br>gcctgatgaatgctcatccggaatttcgtatggcaatgaaagacggtgagctggtgatatgggatagtgttcaccc<br>ttgttacaccgttttccatgagcaaactgaaacgttttcatcgctctggagtgaataccacgacgatttccggcagttt<br>ctacacatatattcgcaagatgtggcgtgttacggtgaaaacctggcctatttccctaaagggtttattgagaatatgt |

| | |
|---|---|
| | ttttcgtctcagccaatccctgggtgagtttcaccagttttgatttaaacgtggccaatatggacaacttcttcgcccc<br>cgttttcaccatgggcaaatattatacgcaaggcgacaaggtgctgatgccgctggcgattcaggttcatcatgcc<br>gtttgtgatggcttccatgtcggcagaatgcttaatgaattacaacagtactgcgatgagtggcagggcggggcgt<br>aatttgatatcgagctcgcttggactcctgttgatagatccagtaatgacctcagaactccatctggatttgttcagaa<br>cgctcggttgccgccgggcgtttttattggtgagaatccaagcctccgatcaacgtctcattttcgccaaaagttgg<br>cccagggcttcccggtatcaacagggacaccaggatttatttattctgcgaagtgatcttccgtcacaggtatttatt<br>cggcgcaaagtgcgtcgggtgatgctgccaacttactgatttagtgtatgatggtgtttttgaggtgctccagtggct<br>tctgtttctatcagctgtccctcctgttcagctactgacggggtggtgcgtaacggcaaaagcaccgccggacatc<br>agcgctagcggagtgtatactggcttactatgttggcactgatgagggtgtcagtgaagtgcttcatgtggcagga<br>gaaaaaaggctgcaccggtgcgtcagcagaatatgtgatacaggatatattccgcttcctcgctcactgactcgct<br>acgctcggtcgttcgactgcggcgagcggaaatggcttacgaacggggc |

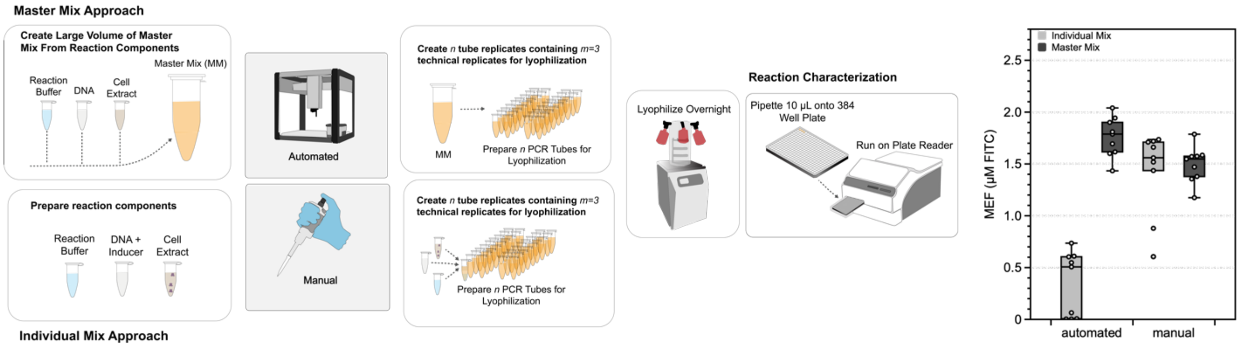


**Supplemental Figure 1**. Individual vs. master mix approaches for robotic reaction construction for pJ23119-sfGFP expression. Schematic shows how individual mixtures and master mixes are prepared, where in the individual mix approach each reaction component is added to tubes serially to complete the reaction. For the master mix approach, a bulk reaction is created and distributed into reaction vessels. The data shows the performance of default Opentrons OT-2 settings and manually prepared reactions under both regimes. Each box and whisker plot represents the end point fluorescence after 8 hours for n=9 reactions.

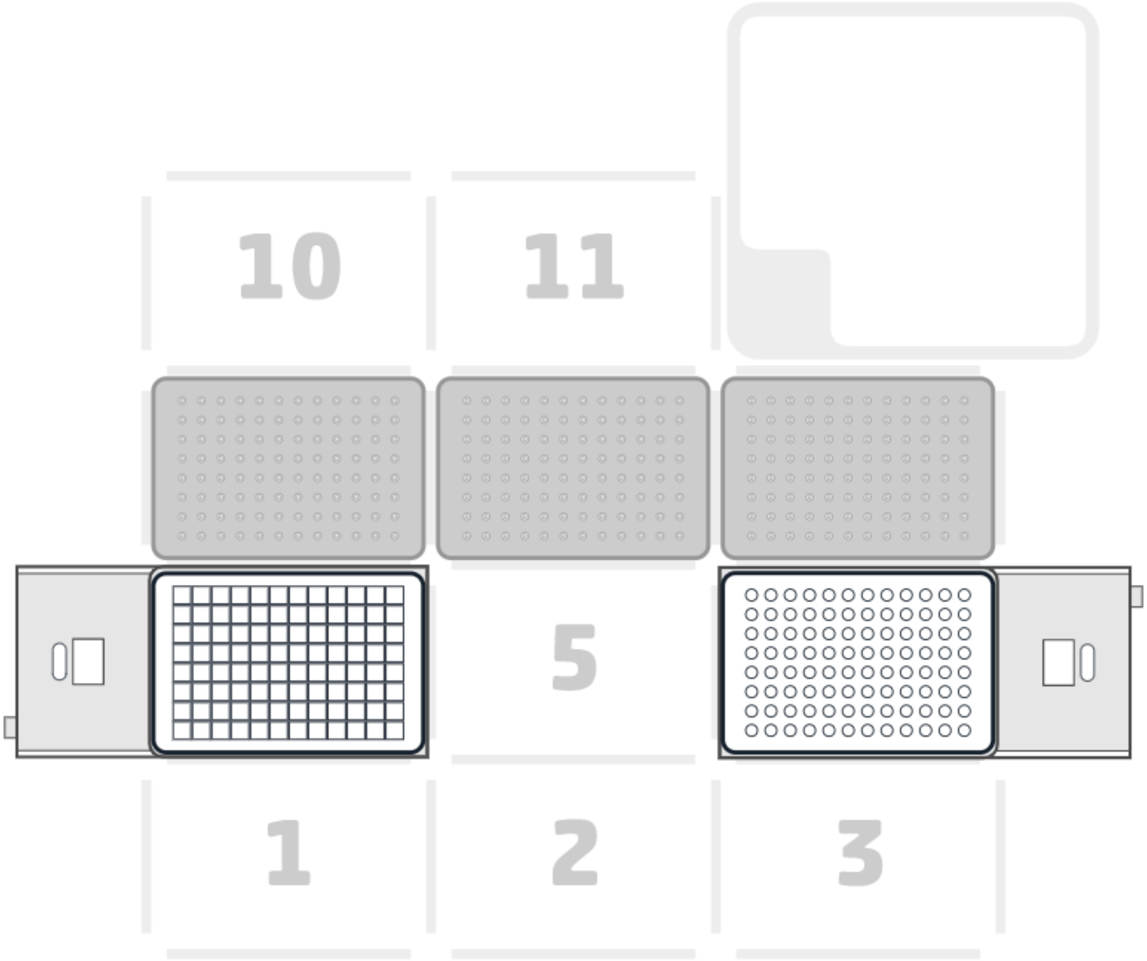


**Supplemental Figure 2**. Opentrons OT-2 deck view for construction of cell-free reactions. Materials required include an Opentrons p20 Multichannel attachment, two Opentrons temperature modules, 3 Opentrons p20 pipette tips, an Opentrons 96-well PCR tube aluminum block, and a 96 deep-well plate.

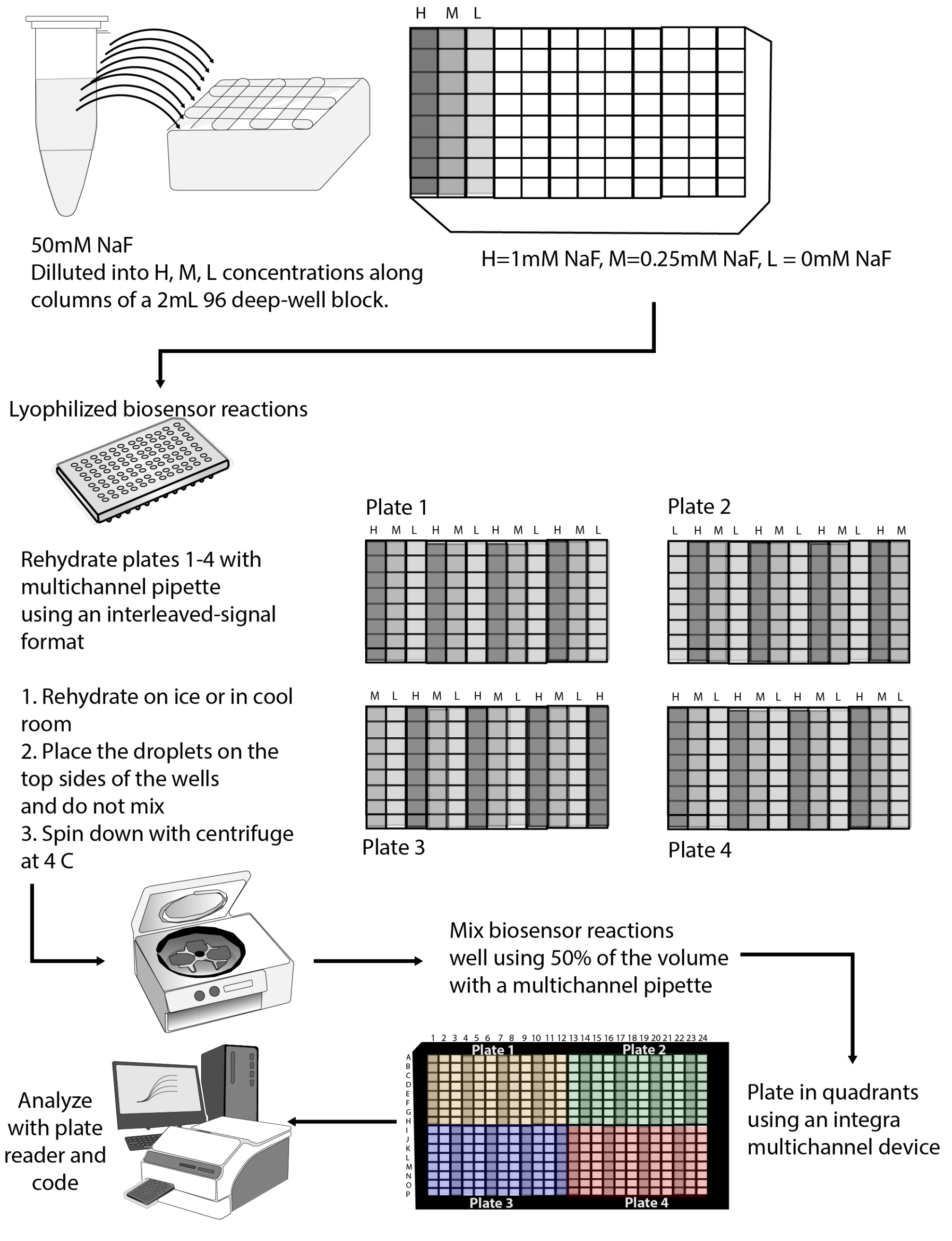


**Supplemental Figure 3**. Approach for rehydrating and analyzing lyophilized reactions using an interleaved signal format for high quality reaction function. Solutions of 1mM (High = H), 0.25mM (Medium = M), and 0mM (Low = L) NaF are made from 50mM NaF stock into the first three rows of 96 deep-well plate. The lyophilized biosensor plates are then rehydrated with the indicated solution using a multichannel pipet on ice, spun down, and mixed using multichannel pipettes. The reactions are then placed into quadrants of a 384 well plate and analyzed on a plate reader at 30°C using emission/excitation 485/520 with fluorescence intervals taken every 5 minutes for 10 hours.
